# Normal and Aberrant Cell Cycles Characterized by a Continuous-Time Stochastic Boolean Model of Cell Cycle Regulation in Budding Yeast

**DOI:** 10.1101/2024.01.04.574212

**Authors:** Kittisak Taoma, John J. Tyson, Teeraphan Laomettachit, Pavel Kraikivski

## Abstract

The cell cycle of budding yeast is governed by an intricate protein regulatory network whose dysregulation can lead to lethal mistakes or aberrant cell division cycles. In this work, we model this network in a Boolean framework for stochastic simulations. Our model is sufficiently detailed to account for the phenotypes of 41 mutant yeast strains (85% of the experimentally characterized strains that we simulated) and also to simulate an endoreplicating strain (multiple rounds of DNA synthesis without mitosis) and a strain that exhibits ‘Cdc14 endocycles’ (periodic transitions between metaphase and anaphase). Because our model successfully replicates the observed properties of both wild-type yeast cells and many mutant strains, it provides a reasonable, validated starting point for more comprehensive stochastic-Boolean models of cell cycle controls. Such models may provide a better understanding of cell cycle anomalies in budding yeast and ultimately in mammalian cells.

## Introduction

Orderly progression through the eukaryotic cell cycle is governed by molecular circuits that control the timely switching from G_1_ into S-G_2_-M and back to G_1_. These transitions typically follow one another in an alternating sequence, but certain disruptions of the control circuits can result in aberrant cell cycles. For example, G_1_-S-G_1_-S and M-(G_1_)-M-(G_1_) cycles are observed in some budding yeast mutant strains (Lu & Cross, 2010; Manzoni et al., 2010; Simmons Kovacs et al., 2012). Moreover, aberrant cell divisions are common occurrences in cancer cells (Edgar et al., 2014; Shu et al., 2018).

Ordinary differential equations (ODEs) are often used to model the molecular control circuits governing cell cycle progression and to explain the irreversible transitions from one cell cycle phase to the next. ODEs have been successfully applied to the complex cell cycle regulatory network in budding yeast (Chen et al., 2000, 2004; Kraikivski et al., 2015), as well as specific cell cycle transitions controlled by different checkpoints, e.g., the G_1_/S transition (Barberis et al., 2007), mitotic exit (Hancioglu & Tyson, 2012; Vinod et al., 2011) and the spindle positioning checkpoint (SPOC) (Howell et al., 2020). Although ODE-based approaches can provide comprehensive quantitative details, they require accurate estimation of many kinetic parameters in the equations and substantial computational time to simulate large molecular regulatory networks (Karlebach & Shamir, 2008). Furthermore, accounting for stochastic effects within this framework requires additional quantitative data about cell constituents and significantly greater computational resources (Barik et al., 2016; Tyson et al., 2019).

To address these difficulties with ODE modeling, many authors have turned to Boolean methods (Davidich & Bornholdt, 2013; Fauré et al., 2009; Li et al., 2004). Recently we have adopted a Boolean Kinetic Monte Carlo (BKMC) approach (Stoll et al., 2012) to explore stochastic Boolean modeling of the budding yeast cell cycle (Laomettachit et al., 2022). Although simple (only seven regulatory proteins), the model successfully explained some basic observations of stochastic cell growth and division in wild-type yeast strains; but it was too simple to account for the phenotypes of any mutant strains. Our goal here is to develop a more comprehensive model that is consistent with certain well-characterized mutant phenotypes, including aberrant cycles such as endoreplication and Cdc14 endocycles (Novak & Tyson, 2022).

The molecular mechanism of our model (Figure 1) involves 22 cell-cycle related components: fifteen proteins, three checkpoints, three ‘progress’ variables, and one ‘flag’.

**Figure 1:**
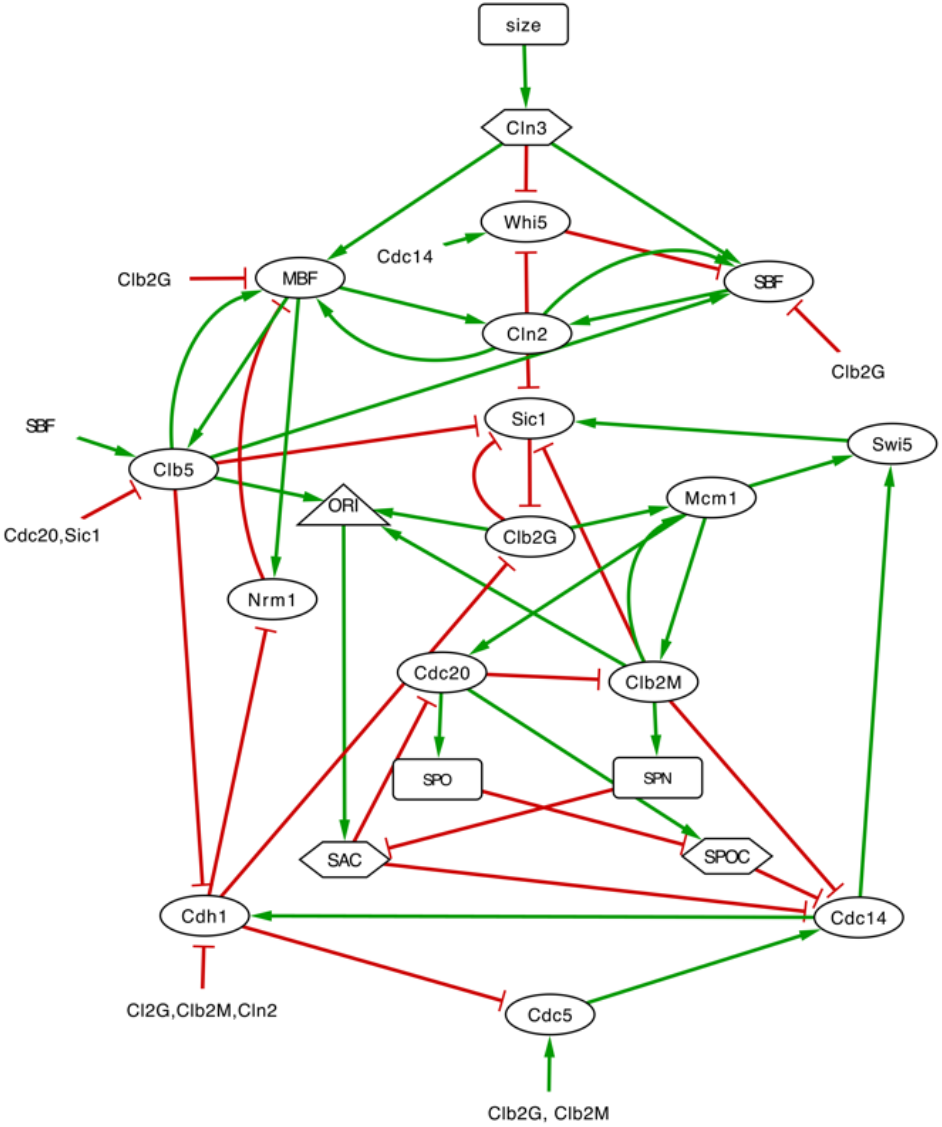
Influence diagram describing our model of cell cycle regulation in budding yeast. The regulatory network consists of nodes connected by edges representing inhibition (red lines with a blunt end) or activation (green lines with a barbed end). Ovals: proteins; hexagons: checkpoints; rectangles: progress variables; triangle: flag variable.

## Method

### Definition of the model

Our model is based on previous work (Stoll et al., 2012; Laomettachit et al., 2022), where the Boolean functions and time steps are updated asynchronously and continuously using Gillespie’s stochastic simulation algorithm. We have extended and modified our earlier model in order to account for the phenotypes of mutant strains as well as the physiology of wild-type cells. Each protein in the model is characterized by a Boolean variable, *X*_*j,t*_, where *j* = 1, …, 15 indexes the proteins and *t* ≥ 0 is time. The Boolean variables are updated by a uniform asynchronous scheme. First, we determine which variables might possibly change in the next step of the Boolean algorithm:

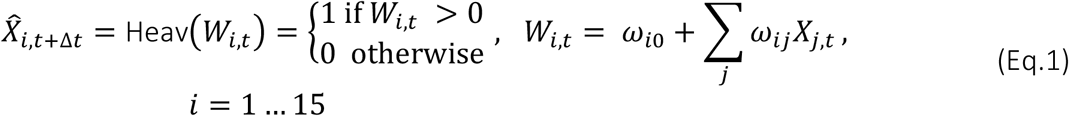

In Eq. 1 the ‘hat’ indicates the ‘potential’ value of *X*_*i*_ at time *t*+Δ*t*. Equation 1 takes as input the 15 Boolean variables representing the proteins and the 3 Boolean variables representing the state of the checkpoints in order to calculate an intermediate function, *W*_*i*_(*X*_1_, …, *X*_15_, *Cln3, SAC, SPOC*), and outputs a Boolean variable, 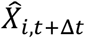, the potential update of *X*_*i*_. The *ω*_*ij*_’s define the Boolean function for updating *X*_*i*_. For example, the Boolean function *X*_1_ = *X*_2_ AND (NOT*X*_3_) can be implemented by the function *W*_1_ = −0.5 + *X*_2_ – *X*_3_; and *X*_1_ = *X*_2_ OR (NOT *X*_3_) by *W*_1_ = +0.5 + *X*_2_ – *X*_3_. In general, *ω*_*ij*_ > 0 if variable *j* activates variable *i* and *ω*_*ij*_< 0 if variable *j* inhibits variable *i*. The *ω*_*ij*_’s are pure numbers, not rate constants. Their values, being fixed by the Boolean relations that define the control system, are not adjusted to fit experimental observations. They are changed, however, to model mutant strains, as explained below.

The *W*_*i*_ functions that define our Boolean model for the 15 cell-cycle regulatory proteins are displayed in Table 1. The 63 *ω*_*i*_ and *ω*_*ij*_ coefficients in these functions are chosen once-and-for-all to fix the logical relations in the Boolean dynamics. Their values (for wild-type cells) are specified in Supplementary Table S1A.

**Table 1:** The *W*_*i*_ functions defining the Boolean network for updating the 15 proteins of the cell-cycle control system. The three checkpoint variables are highlighted in red.

|  |  |
| --- | --- |
| 1 | $W_{\text{Whi5}} = \omega_{\text{Whi5}} + \omega_{\text{Whi5,Cdc14}} \cdot \text{Cdc14} - \omega_{\text{Whi5,Cln2}} \cdot \text{Cln2} - \omega_{\text{Whi5,Cln3}} \cdot \text{Cln3}$ |
| 2 | $W_{\text{SBF}} = \omega_{\text{SBF}} - \omega_{\text{SBF,Whi5}} \cdot \text{Whi5} - \omega_{\text{SBF,Clb2G}} \cdot \text{Clb2G} + \omega_{\text{SBF,Clb5}} \cdot \text{Clb5} + \omega_{\text{SBF,Cln2}} \cdot \text{Cln2} + \omega_{\text{SBF,Cln3}} \cdot \text{Cln3}$ |
| 3 | $W_{\text{MBF}} = \omega_{\text{MBF}} - \omega_{\text{MBF,Nrm1}} \cdot \text{Nrm1} - \omega_{\text{MBF,Clb2G}} \cdot \text{Clb2G} + \omega_{\text{MBF,Clb5}} \cdot \text{Clb5} + \omega_{\text{MBF,Cln2}} \cdot \text{Cln2} + \omega_{\text{MBF,Cln3}} \cdot \text{Cln3}$ |
| 4 | $W_{\text{Nrm1}} = \omega_{\text{Nrm1}} - \omega_{\text{Nrm1,Cdh1}} \cdot \text{Cdh1} + \omega_{\text{Nrm1,MBF}} \cdot \text{MBF}$ |
| 5 | $W_{\text{Clb5}} = \omega_{\text{Clb5}} - \omega_{\text{Clb5,Sic1}} \cdot \text{Sic1} - \omega_{\text{Clb5,Cdc20}} \cdot \text{Cdc20} + \omega_{\text{Clb5,MBF}} \cdot \text{MBF} + \omega_{\text{Clb5,SBF}} \cdot \text{SBF}$ |
| 6 | $W_{\text{Cln2}} = \omega_{\text{Cln2}} + \omega_{\text{Cln2,SBF}} \cdot \text{SBF} + \omega_{\text{Cln2,MBF}} \cdot \text{MBF}$ |
| 7 | $W_{\text{Sic1}} = \omega_{\text{Sic1}} + \omega_{\text{Sic1,Swi5}} \cdot \text{Swi5} - \omega_{\text{Sic1,Clb2G}} \cdot \text{Clb2G} - \omega_{\text{Sic1,Clb2M}} \cdot \text{Clb2M} - \omega_{\text{Sic1,Clb5}} \cdot \text{Clb5} - \omega_{\text{Sic1,Cln2}} \cdot \text{Cln2}$ |
| 8 | $W_{Cdh1} = \omega_{Cdh1} + \omega_{Cdh1,Cdc14} \cdot Cdc14 - \omega_{Cdh1,Clb2G} \cdot Clb2G - \omega_{Cdh1,Clb2M} \cdot Clb2M$ $- \omega_{Cdh1,Clb5} \cdot Clb5 - \omega_{Cdh1,Cln2} \cdot Cln2$ |
| 9 | $W_{Clb2G} = \omega_{Clb2G} - \omega_{Clb2G,Cdh1} \cdot Cdh1 - \omega_{Clb2G,Sic1} \cdot Sic1$ |
| 10 | $W_{Clb2M} = \omega_{Clb2M} + \omega_{Clb2M,Mcm1} \cdot Mcm1 - \omega_{Clb2M,Cdc20} \cdot Cdc20$ |
| 11 | $W_{Mcm1} = \omega_{Mcm1} + \omega_{Mcm1,Clb2G} \cdot Clb2G + \omega_{Mcm1,Clb2M} \cdot Clb2M$ |
| 12 | $W_{Cdc5} = \omega_{Cdc5} + \omega_{Cdc5,Clb2G} \cdot Clb2G + \omega_{Cdc5,Clb2M} \cdot Clb2M - \omega_{Cdc5,Cdh1} \cdot Cdh1$ |
| 13 | $W_{Cdc20} = \omega_{Cdc20} + \omega_{Cdc20,Mcm1} \cdot Mcm1 - \omega_{Cdc20,SAC} \cdot SAC$ |
| 14 | $W_{Cdc14} = \omega_{Cdc14} + \omega_{Cdc14,Cdc5} \cdot Cdc5 - \omega_{Cdc14,Clb2M} \cdot Clb2M - \omega_{Cdc14,SAC} \cdot SAC$ $- \omega_{Cdc14,SPOC} \cdot SPOC$ |
| 15 | $W_{Swi5} = \omega_{Swi5} + \omega_{Swi5,Cdc14} \cdot Cdc14 + \omega_{Swi5,Mcm1} \cdot Mcm1$ |

If more than one protein potentially changes state 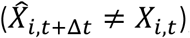, then we choose the one that will actually change at random with equal probabilities, i.e., the Boolean model progresses from one state to the next by uniform asynchronous updating. In the BKMC approach, time is updated by choosing Δ*t* from an exponential distribution parameterized by the total ‘propensity’ (probability per unit time) for any one of the potential changes to occur. This scheme is based on the assumption that each potential change is an elementary chemical reaction (Gillespie, 2007), which certainly doesn’t hold in the case of Boolean modeling. Since each change (protein synthesis, degradation, phosphorylation, dephosphorylation, etc.) is a complex series of elementary reactions, we compute the time increment, Δ*t*, at which the selected change takes place from a gamma distribution, with density function:

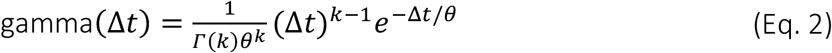

The parameters *k* and *θ* determine the mean value of Δ*t* between updates (mean = *kθ*) and its coefficient of variation 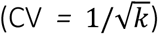. For proteins that change slowly (e.g., by synthesis and degradation), we set *k* = 3 and *θ* = 0.3 min (i.e., mean = 0.9 min and CV *=* 0.58); and for proteins that change more quickly (e.g., by phosphorylation and dephosphorylation), we set *k* = 2 and *θ* = min (i.e., mean = 0.6 min and CV *=* 0.71). See Supplementary Table S2 for our classification of proteins as slow or fast. In some circumstances, the Boolean model of the protein interactions settles on a steady state (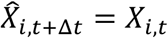 for all *i* = 1, …, 15), in which case we allow *t* to increase by drawing Δ*t* from a gamma distribution with *k* = 5 and *θ* = 0.3 min (i.e., mean = 1.5 min and CV *=* 0.45). During the ensuing period, the ‘progress’ variables may change and induce the protein network to leave the steady state and re-enter the cell cycle.

For instance, wild-type cells have a G_1_ steady state (*Whi5* = *Cdh1* = *Sic1* = 1, all other Boolean variables = 0). As formalized below, for a cell in this G_1_ steady state, the progress variable *size*_*t*_ steadily increases as the cell grows. When *size*_*t*_ > *S*_0_, the ‘cell size checkpoint’ is satisfied, and the checkpoint variable *Cln3*_*t*_ is changed from 0 to 1. This change kicks the protein interaction network out of the G_1_ steady state and sets the cell division program in motion. In the simulation of some mutants, the protein interaction network falls into a steady state that it cannot leave (i.e., the cell is arrested at some point in the cell cycle), and we stop the simulation after the arrested state becomes evident.

After the updated protein (say, *X*_*k*_) is chosen and Δ*t* is determined, all the protein variables are updated as follows:

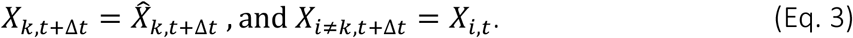

In addition to the Boolean variables tracking the protein interaction network, the model has a Boolean ‘flag’ called ORI and three Boolean ‘checkpoints’ called Cln3, SAC and SPOC.

- **ORI** specifies the state of the origins of replication on the chromosomes. *ORI* = 0 means the chromosomes are unreplicated and the origins are ‘licensed’ to initiate replication. *ORI* = 1 means that chromosome replication has been initiated and that the origins are now ‘unlicensed’ (i.e., unable to initiate a new round of DNA replication). The value of *ORI* at any time *t* is determined simply by the presence of Clb-dependent kinase activity:

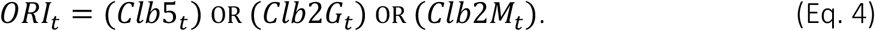
- **Cln3** is an indicator of cell growth. *Cln3* = 0 indicates that a cell is too small to start S phase; *Cln3* = 1 means that it has grown large enough to warrant a new round of DNA replication and cell division:

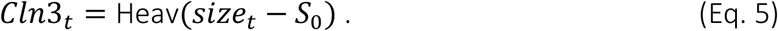

‘Size’ is the progress variable that controls the size checkpoint *size*_*t*_ > *S*_0_, where *S*_0_ is the minimum size necessary start the S-G2-M sequence. *S*_0_ is a random variable assigned to a cell at birth. *S*_0_ is chosen from a lognormal distribution,

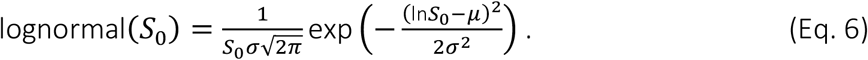

For a lognormal distribution, 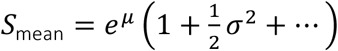 and 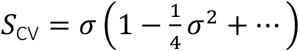. For simplicity, we encode the lognormal distribution with *μ* = ln(*S*_0_mean_) and *σ* = *S*_0_CV_, which are suitable approximations for our purposes.

During every time step Δ*t, size*_*t*_ increases according to:

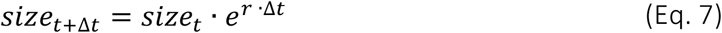

where *r* is the specific growth rate of cells. At cell division (an event to be defined later), the size of the dividing cell, *size*_@div_, is distributed asymmetrically to the newborn cells:

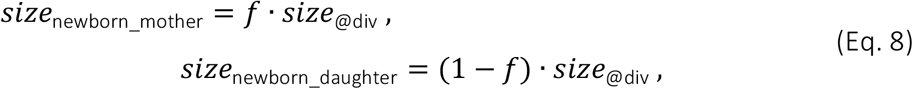

where *f* is drawn from a lognormal distribution with *μ* = ln(*f*_mean_) > 0.5 and *σ* = *f*_CV_.

- **SAC**, the ‘spindle assembly checkpoint’, is a Boolean variable indicating the state of alignment of replicated chromosomes on the mitotic spindle:

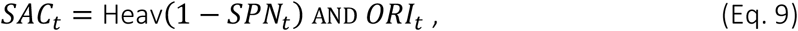

where *ORI*_*t*_ = 1 indicates that DNA replication has been initiated, and *SPN*_*t*_ tracks the progression of the replicated chromosomes on the spindle; *SPN*_*t*_ = 1 indicating complete alignment. *SPN*_*t*_ is initialized at 0 when the cell enters mitosis, i.e., when Clb2M turns on, and *SPN*_*t*_ increases in each time step thereafter, according to:

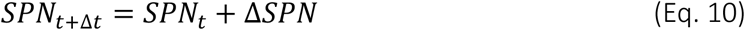

where Δ*SPN* is chosen (in each time step) from a lognormal distribution with parameters *SPN*_mean_ and *SPN*_CV_.
- **SPOC**, the ‘spindle position checkpoint’, is a Boolean variable indicating that the fully aligned mitotic spindle is properly positioned in the neck between mother and bud:

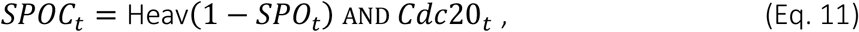

where *Cdc20*_*t*_ = 1 indicates that anaphase has been initiated, and *SPO*_*t*_ tracks the movement of the two incipient nuclei during anaphase and telophase. *SPO*_*t*_ = 1 indicates that the bud has received its nucleus. When SPOC turns off, Cdc14 is activated and the cell completes the transition from telophase to G_1_. *SPO*_*t*_ is initialized at 0 when the cell enters anaphase, i.e., when Cdc20 turns on, and *SPO*_*t*_ increases in each time step thereafter, according to:

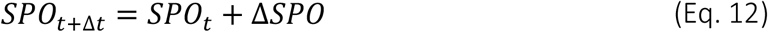

where Δ*SPO* is chosen (in each time step) from a lognormal distribution with parameters *SPO*_mean_ and *SPO*_CV_. In addition to the 63 *ω*_*i*_ and *ω*_*ij*_ coefficients defined in Suppl. Table S1A, the equations defining our model involve 16 adjustable parameters: 4 parameters for updating the SPN and SPO variables, 2 parameters for the specific growth rate (mass doubling time) in glucose and galactose media, 2 parameters to determine the fraction *f* of a dividing cell that is apportioned to the mother cell, and 2 parameters to determine the critical cell size *S*_0_ (see Suppl. Table S1B), and 6 parameters to define Δ*t* for fast and slow protein changes and when no proteins change state (see Suppl. Table S2). These 16 parameters are manually adjusted to fit the model to experimental observations in both wild-type and mutant strains. See the supplementary tables for their values in wild-type cells.

### Experimental evidence for the Boolean model

Cell cycle progression through G_1_ phase is inhibited by Whi5, which is inactivated (phosphorylated) by Cln3, Bck2, Cln1, and Cln2 (de Bruin et al., 2004, Skotheim et al., 2008) and activated (dephosphorylated) by Cdc14 phosphatase (Table 1, Row 1). In our model, the Cln3 variable accounts for both Cln3 and Bck2 proteins, and the Cln2 variable represents both Cln1 and Cln2. Inactivation of Whi5 promotes activation of the SBF transcription factor (Row 2). Also, the activities of both SBF and MBF transcription factors are upregulated by Cln3, Cln2, and Clb5 (Rows 2-3) (Charvin et al., 2010; Dirick & Nasmyth, 1991; Koch et al., 1996; Nasmyth, 1993). Activation of SBF and MBF defines the START transition in the budding yeast cell cycle, after which yeast cells set off on an irreversible path to DNA synthesis, mitosis and cell division. Along this path, SBF and MBF are inactivated by Clb2 cyclin (Rows 2-3). In addition, MBF is regulated by a negative feedback loop with Nrm1 (Rows 3-4) (Ostapenko & Solomon, 2011)

After the START transition, MBF and SBF activate the synthesis of Clb5, Clb6, and Cln1, Cln2 cyclins, which are responsible for DNA replication (Row 5) and budding (Row 6), respectively (Wittenberg & Reed, 2005). (In our notation, the *Clb5* variable represents both Clb5 and Clb6 cyclins.) The origin licensing variable ORI, is set to 0 (origins licensed) when both Clb5 and Clb2 are inactivated as a mother cell exits mitosis and divides, then *ORI* is flipped to 1 (DNA replication begins) when either Clb5 or Clb2 is activated in the next cell cycle (Eq. 4).

Once DNA replication is initiated, Clb5 activity promotes the accumulation of active Clb1 and Clb2 cyclins by suppressing Sic1 (Row 7) and Cdh1 (Row 8) in late S phase (Yeong et al., 2001). Subsequently, a positive feedback loop with Mcm1 transcription factor sets off rapid accumulation of Clb1 and Clb2 cyclins, which drive the cell into M phase (Maher et al., 1995) (Rows 9-11). Clb2G represents Clb1, Clb2 cyclin activities in late S phase and Clb2M represents a higher level of Clb1, Clb2 cyclins in M phase. With the high level of Clb2 (*Clb2M* = 1), the cell enters M phase and Cdc5 is activated (Row 12).

In M phase, the spindle assembly checkpoint (SAC) prevents the metaphase-to-anaphase transition until all sister chromatids achieve bipolar alignment on the mitotic spindle. The SAC turns ON (*SAC* = 0 → 1) when DNA replication begins (*ORI* = 1) (Eq. 9). Progress in aligning the replicated chromosomes on the mitotic spindle is tracked by the SPN variable. When the cell enters M phase (*Clb2M* = 1), the (continuous) variable *SPN*_*t*_ starts to increase (Eq. 10). When *SPN*_*t*_ reaches 1 (i.e., all chromosomes are aligned on the metaphase plate), *SAC* is set to zero.

Once the cell passes the SAC (*SAC* = 1 → 0), Cdc20 is activated (*Cdc20* = 0 → 1) and it promotes the metaphase-anaphase transition. *SPN*_*t*_ is reset to zero, and the Spindle Position Checkpoint (SPOC) is activated (*SPOC* = 0 → 1) to ensure that both mother and daughter cells receive a full set of chromosomes before cytokinesis (Caydasi et al., 2010). Spindle positioning is monitored by the (continuous) SPO variable (Eq. 12). When *SPO*_*t*_ = 1, the SPOC is satisfied (*SPOC* = 1 → 0), and *SPO* is reset to 0.

To exit mitosis, Cdc14 must be fully activated (i.e., released from the RENT complex in the nucleolus), which is a consequence of both the FEAR and MEN pathways (Queralt & Uhlmann, 2008). FEAR is activated when *SAC* → 0 and MEN when *SPOC* → 0 (Row 14). Finally, Cdc14 activates Cdh1 and Swi5 (Rows 8 & 15), and Swi5 (a transcription factor) activates Sic1 (Row 7). Together Sic1 and Cdh1 reverse the activities of all cyclins as the newly divided cells re-enter G_1_ (Visintin et al., 1998).

### Description of mutant simulations

A mutant strain in which ‘*gene K*’ is deleted is modeled by setting *ω*_*k,j*_ = 0 for all coefficients that describe the activation of protein *K*. For example, the *whi5 Δ* mutant strain is modeled by setting ***ω***_**whi5**_ **= 0** and ***ω***_**whi5**,**Cdc14**_**= 0**. Mutant strains containing a GAL promoter of gene *L* are modeled by adding 0.5 to the *ω*_*L,0*_ function in Eq. 1, in order to change the activation threshold for variable *X*_*L*_ (the activity of the protein *L*). For example, *GAL-SIC1* is modeled by adding 0.5 to Table 1 Row 7, to account for a more potent promoter of Sic1 expression in the *GAL-SIC1* strain. Also, the growth rate constant for galactose medium, *r* = 0.0046 min_−1_, is used for all mutant strains with the GAL promoter. For mutant strains that exhibit endoreplication and Cdc14 endocycles, we also change the parameters of the gamma distribution (Eq. 2) in order to match the period of oscillation to experimental observations. For example, for the *GAL-CLB2-db* Δ mutant strain, Clb2 (which inhibits Cdh1) is non-degradable and its level is high, therefore the activation of Cdh1 is delayed compared to wild-type cells with normal levels of Clb2. Also, because Clb2 activates Cdc5, the inactivation of Cdc5 is delayed when the level of Clb2 is high. Therefore, for this mutant strain, the gamma-distribution parameters are set to *μ* = 22.5 min and *CV =* 0.25 for calculating Δ*t* for Cdc5 inactivation and Cdh1 activation. Similar reasoning applies to the *clb1-5*Δ mutant strain that exhibits endoreplication cycles. In the absence of most Clbs, Cdh1 inactivation and Clb6 activation are delayed, and therefore, we set *μ* = 22.5 min and *CV =* 0.25 for the timing of these events in this mutant strain. All time delay parameters that are used to model mutant strains exhibiting endoreplication and Cdc14 endocycles are further explained in Supplementary Table S4.

## Results

To assess the potential of our method, we present simulations of wild-type cell cycles, of population-level properties of budding yeast cultures, and of mutant strains that exhibit aberrant cycles. We also use the model to predict phenotypes of mutant strains that have not yet been characterized experimentally.

### Simulation of cell cycle progression in wild-type cells

Figure 2 shows simulations of key components regulating cell cycle events in wild-type budding yeast. Newborn yeast cells must grow to a ‘critical size’ in order to activate Cln3 and subsequently to inactivate Whi5, which then permits activation of SBF and MBF transcription factors (Fig. 2A). MBF induces the synthesis of Clb5, which induces DNA replication (ORI = 1 identifies the onset of S phase, Fig. 2B). The spindle assembly progress variable (*SPN*) indicates progression through G_2_/M into metaphase. When *SPN* ≥ 1, the spindle assemble checkpoint variable (*SAC*) is set to zero, which allows the activation of Cdc20 and the cell to progress into anaphase (see Figure 2C).

**Figure 2:**
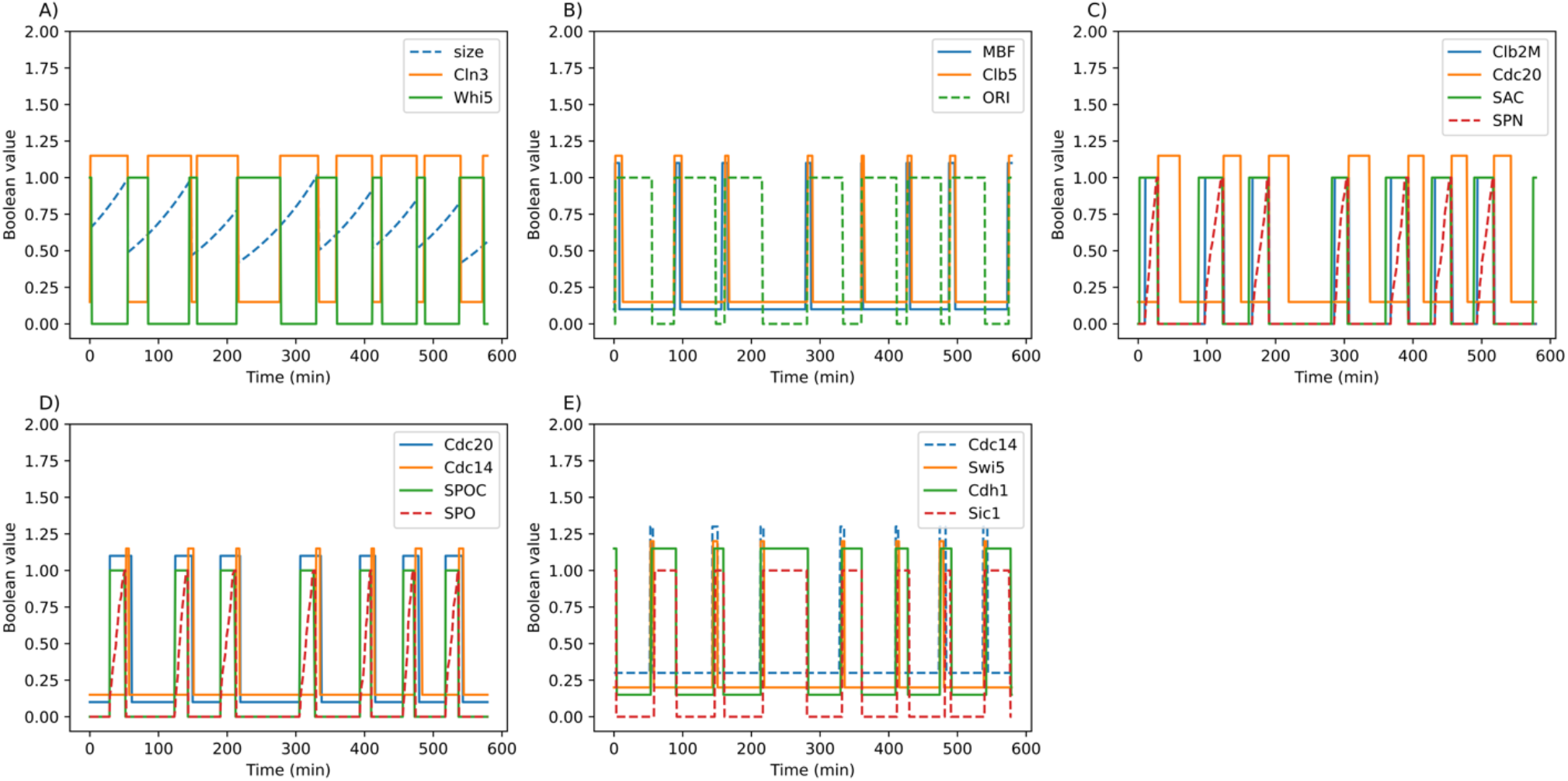
Simulation results for wild-type budding yeast cell cycles. Each subplot shows the dynamics of key cell cycle components for seven division cycles. (A) Cell size and molecular components that regulate the progression through G_1_. (B) The initiation of DNA replication. (C) Spindle assembly progress in response to the activation of Clb2M. (D) Spindle orientation progress after anaphase. (E) Cdc14 activation and resetting into G_1_. Activities of proteins in the panels are offset for clearer visualization.

The spindle-orientation progress variable (*SPO*) accounts for proper segregation of chromosomes into mother and daughter cell compartments during anaphase. When *SPO* ≥ 1, the spindle orientation checkpoint (*SPOC*) is set to zero, which allows Cdc14 to be fully released from the nucleolus (Figure 2D). Once Cdc14 is released, it activates Cdh1 and Swi5 (which initiates synthesis of Sic1), thereby resetting the cell back to G_1_ (Figure 2E).

Next, the model is used to simulate the exponential expansion of a population of budding yeast cells, in order to estimate the means and standard deviations of observable cell-cycle measures: the period from birth to division (*T*_c_), the duration of G_1_ phase from mitotic exit to S phase (*T*_G1_), the period from budding to division (*T*_bud_), and cell size at birth. We compare these simulation results with corresponding experimental data from (Di Talia et al., 2007) in Figure 3. Overall, the model accurately simulates these population-level properties in both mother cells (Figure 3A and B) and daughter cells (Figure 3D and E); although, the model overestimates *T*_G1_ variability in both mother and daughter cells. Similar discrepancies were observed in simulations based on previous models (Laomettachit et al., 2016, 2022). Figures 3C and F show the joint distributions of size-at-birth and *T*_G1_ for mother and daughter cells. The simulated data points are fitted with trendlines, as was done to analyze the experimental data (Di Talia et al., 2007). The estimated slopes of the trendlines for mother cells (−0.65) and daughter cells (large −0.58 & small −0.99) show similar trends to the experimental data (slope = −0.1 for mother cells; −0.3 & −0.7 for large & small daughter cells, respectively).

**Figure 3:**
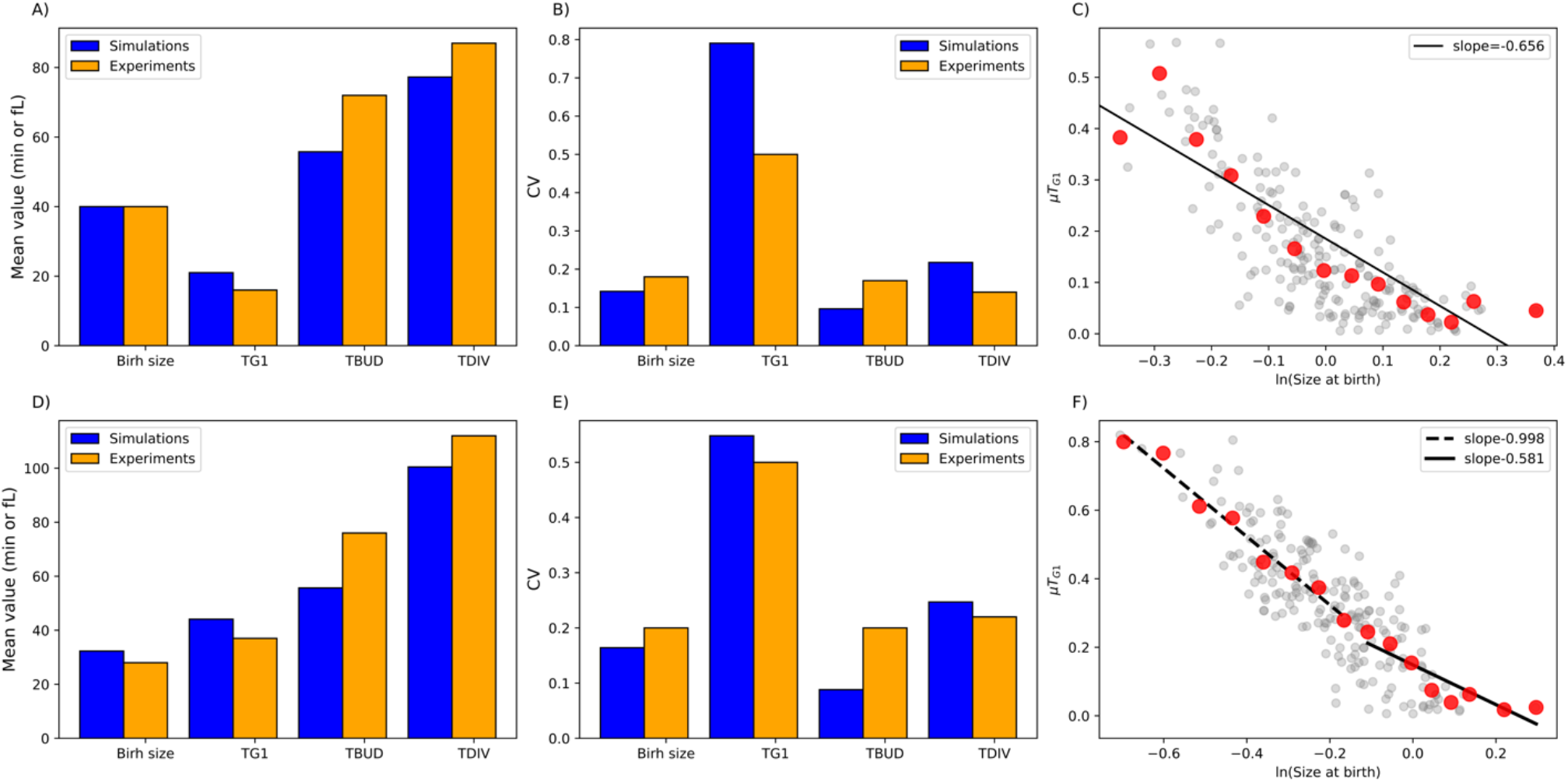
Population-level statistics from model simulations and experimental observations. The mean values and coefficients of variation for four cell-cycle properties in populations of mother cells (A and B) and daughter cells (D and E). To transform dimensionless cell size into volume in fL, we used a conversion factor of 74.66 fL, which was derived by equating the mean size of mother cells to the experimental mean volume of ∼40 fL. To visualize the joint distributions of size at birth and G_1_ duration (*T*_G1_) in mother cells (C) and daughter cells (F), we plot (grey dots) 200 simulated cells sampled from the whole population. To estimate the trends in the data, we plot (red dots) the average value of the grey dots in bins of size 2 fL, exactly as implemented by the authors of the experimental data (Di Talia et al., 2007).

To estimate how fast a simulated population of cells loses synchrony over time, 100 cells with average size of 0.65 were initiated in the G_1_ state and tracked (both mother and daughter cells) for 500-time steps (Figure 4A). Cell size and protein activities of all cells extant at time *t* were averaged and the results plotted as functions of *t* (Figure 4B). The population-average results clearly show a loss of synchrony that agrees well with observations (Woldringh et al., 1993). Protein activities and cell size quickly lose synchrony due to the unequal division of material between daughter and mother cells.

**Figure 4:**
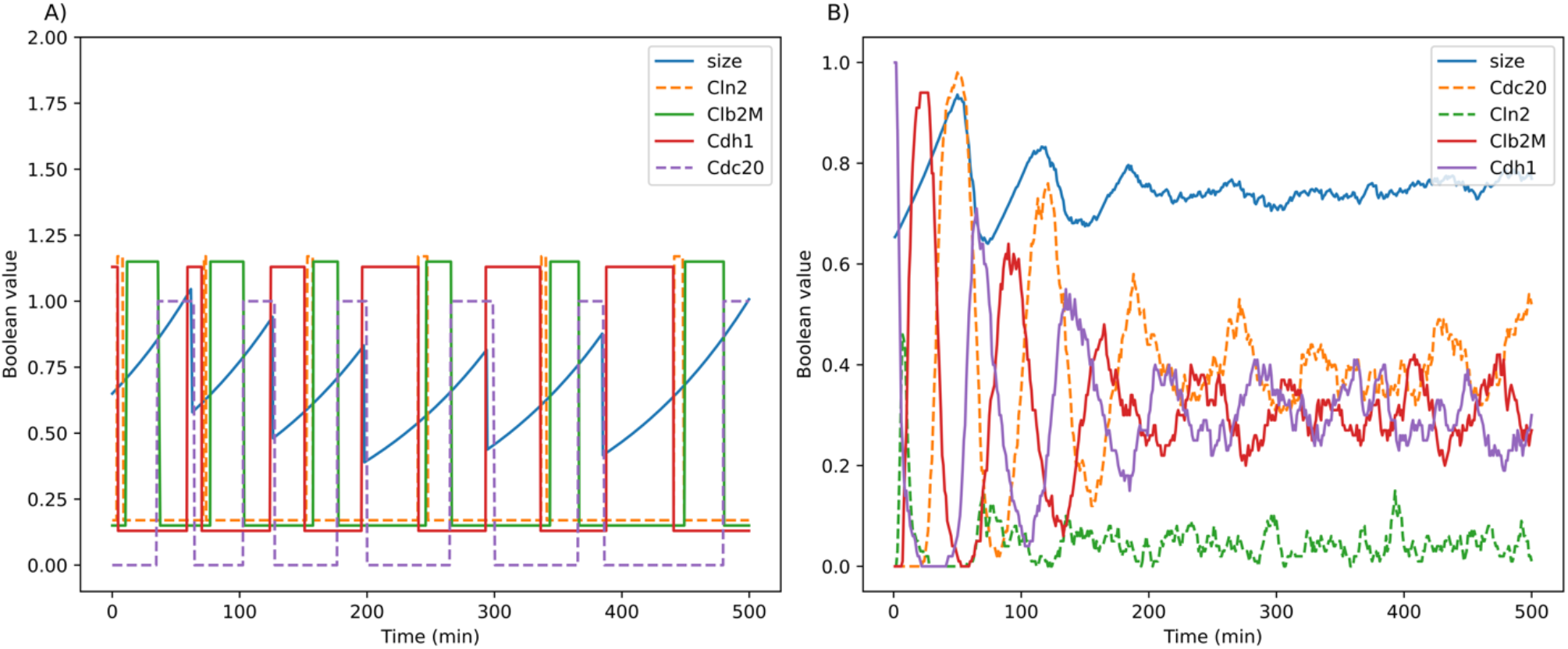
Loss of synchrony of protein activities from model simulations. (A) A single cell is tracked over time from one division to the next. (B) Many such simulations are averaged to predict the loss of synchrony in a population of cells. Activities of proteins in the panel (A) are offset for clearer visualization.

### Simulation of mutant phenotypes

To test the accuracy of our model in accounting for mutant phenotypes, we simulated cell cycle progression in 48 experimentally characterized mutant strains, including gene deletion and overexpression mutants. Of these strains, our model agrees with 41 observed phenotypes. Here we focus on mutant strains exhibiting aberrant cycles. Results for the remaining strains are recorded in Supplementary Table S3.

For the *clb1-5*Δ strain, in which all Clbs—except Clb6—are deleted, cells replicate the genome multiple times without mitosis (Simmons Kovacs et al., 2012), a phenotype called endoreplication. Because the Clb5 variable represents both Clb5 and Clb6, the action of Clb6 in *clb1-5*Δ mutant is simulated by reducing the basal parameter *ω*_Clb5,_ to −1 and the values of parameters ω_*i*,Clb5_ that describe the influence of Clb5 and Clb6 on target protein *i* were reduced by eight-fold (see Supplementary Table S3). Also, due to absence of most Clbs in the *clb1-5*Δ mutant, Cdh1 inactivation is delayed by 22.5 minutes compared to 0.6 minutes in wild type. As Figure 5A shows, the *clb1-5*Δ mutant fails to enter mitosis and to divide, and cell size becomes extremely large after *t* = 250 min when all Clbs except Clb6 are deleted. However, Clb6, Cdh1 and Nrm1 continue to oscillate, driven by the negative feedback loop MBF → Clb6 –| Cdh1 –| Nrm1 –| MBF. Figure 5B shows the distribution of endoreplication periods. The estimated period of Clb6 oscillations, 59.60 ± 0.24 min, is in good agreement with experimental observations (Simmons Kovacs et al., 2012).

**Figure 5:**
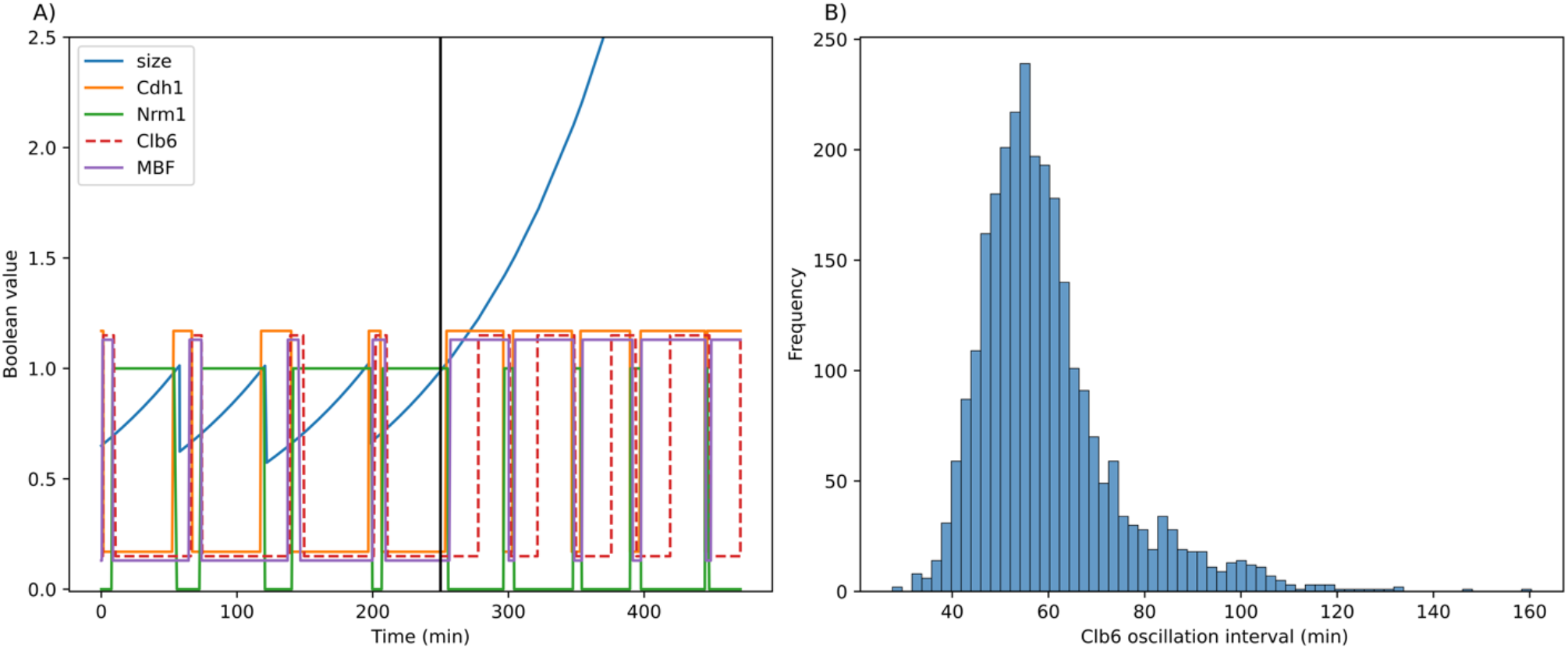
Simulation of endoreplication cycles (A) and the distribution Clb6 oscillation periods (B). Activities of proteins in the panel (A) are offset for clearer visualization.

Endoreplication cycles in our model disappear when any component (i.e., Cdh1, Nrm1 or MBF) in the negative feedback loop is deleted. Further, we tested all pairwise deletions of model components in *clb1-5*Δ mutant strain to identify other mutant strains that might exhibit endoreplication cycles. Mutant strains that lose or retain endoreplication cycles are shown in Supplementary Figure S1.

Another mutant strain that exhibits aberrant cycles is *GAL-CLB2-db*Δ. The high level of non-degradable Clb2 causes cell cycle arrest in mitosis. Although the cells cannot exit mitosis, the high level of Clb2 supports the activation of Cdc5 which promotes Cdc14 release, Cdc14 then activates Cdh1 which degrades Cdc5. This negative loop (Cdc5 → Cdc14 → Cdh1 –| Cdc5) results in Cdc14 endocycles (Figure 6A). Figure 6B shows the distribution of endocycle periods. The averaged period of Cdc14 oscillations is 59.10 ± 0.25 min, in agreement with experimental observations (Lu & Cross, 2010; Manzoni et al., 2010). Other mutant strains with sustained and disrupted Cdc14 endocycles are shown in Supplementary Figure S2.

**Figure 6:**
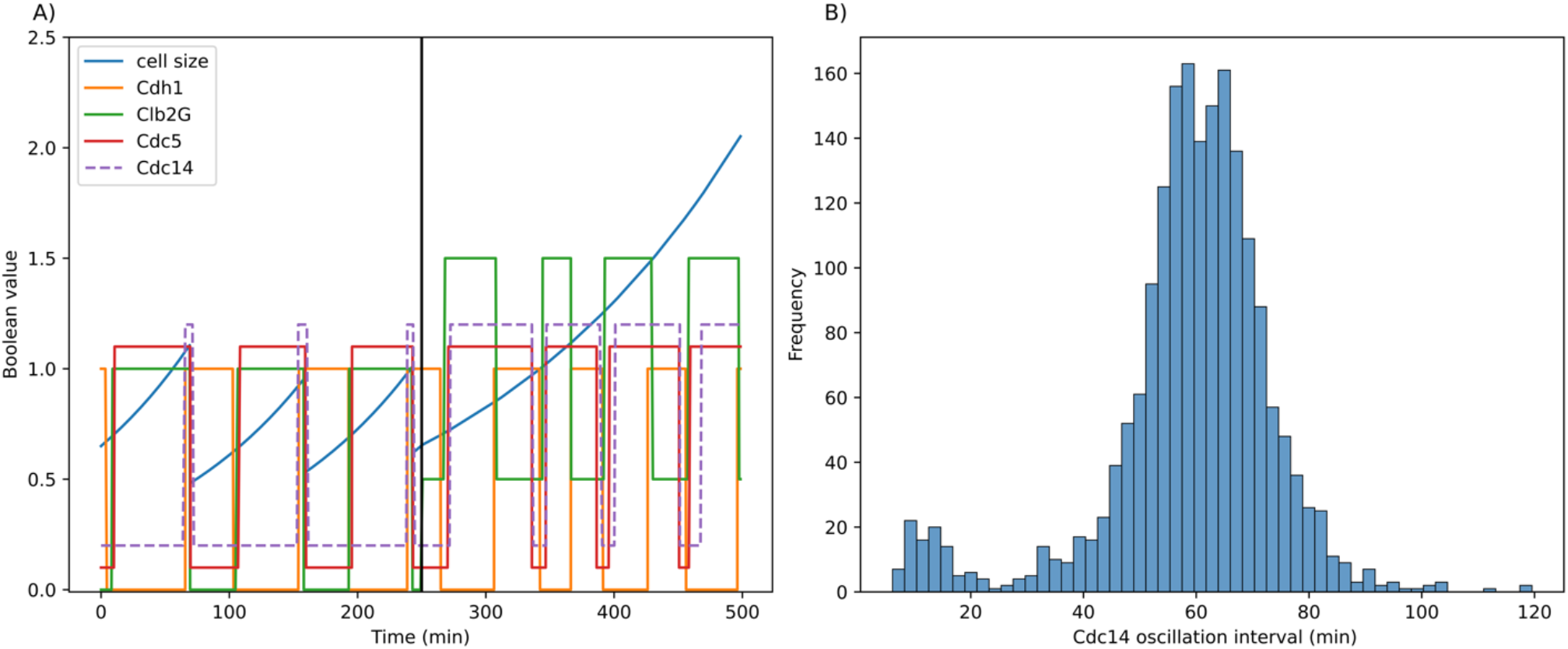
The simulation of Cdc14 endocycles (A) and the distribution of Cdc14 oscillation periods (B). Activities of proteins in the panel (A) are offset for clearer visualization

### Model predictions

Next, we used the model to predict the phenotypes of mutant strains with single and double deletions of model variables. Because some model variables already represent two genes (MBF = SWI6 + MBP1, SBF = SWI6 + SWI4, CLN3 = CLN3 + BCK2, CLN2 = CLN1 + CLN2, CLB5 = CLB5 + CLB6, and CLB2 = CLB1 + CLB2), combining their deletions with the deletion of other model components generates triple- or quadruple-deletion strains of budding yeast. The model, with 15 genetic components, predicts the phenotypes of 15 + 15x14/2 = 120 ‘single’ and ‘double’ deletion strains, among which only 24 have been experimentally characterized. As shown in Figure 7, the model successfully accounts for the phenotypes of 23 known mutants; only *cln3*Δ *bck2*Δ *whi5*Δ, disagrees with its observed phenotype. Ninety-six mutants are not yet experimentally characterized.

**Figure 7:**
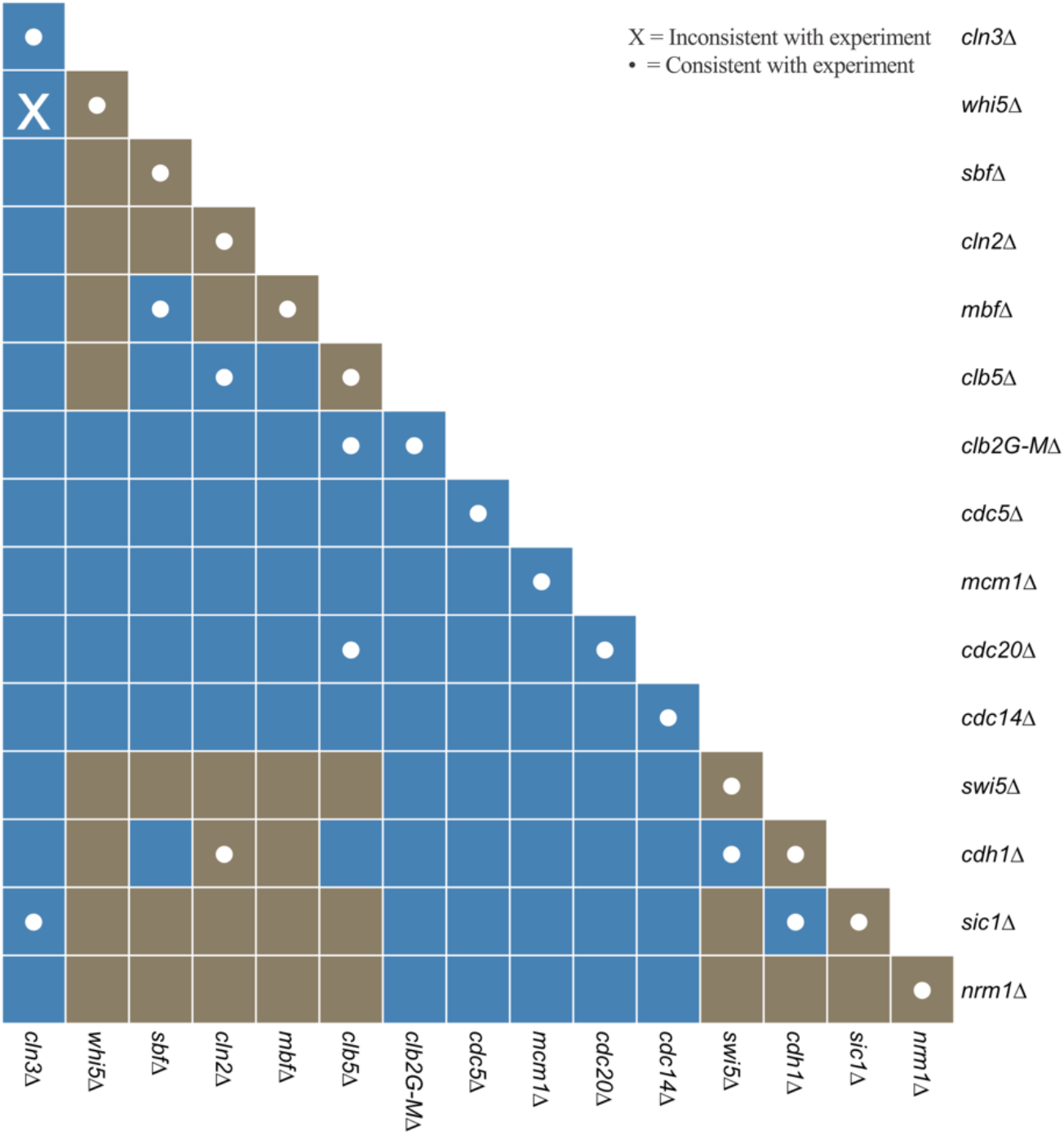
Predicted phenotypes of double-deletion strains. Each rectangular cell corresponds to a combination of two deleted components listed along the vertical and horizontal axes. Blue color represents synthetic lethality and brown color indicates a viable phenotype. A white dot indicates that the prediction is consistent with experimental data, and “X” indicates a difference between the simulated and observed phenotypes. The elements along the diagonal correspond to single-deletion strains.

## Discussion

In this study, we developed a stochastic Boolean model of the budding yeast cell cycle that can correctly explain both normal cell cycle progression and aberrant cycles (endoreplication and Cdc14 endocycles). Understanding the mechanism of aberrant cycles and identifying controls that suppress these cycles in wild-type cells is important because these cycles are found in many types of cancer (Zhang et al., 2022). Our model confirms that endoreplication cycles are driven by the negative feedback loop MBF → Clb6 –| Cdh1 –| Nrm1 –| MBF, and Cdc14 endocycles are driven by the negative loop Cdc5 → Cdc14 → Cdh1 –| Cdc5. Our model also successfully accounts for the phenotypes of 41 gene deletion and overexpression mutant strains out of 48 strains tested. Furthermore, we used the model to predict the phenotypes of 120 mutant strains carrying one-, two-, three- or four gene deletions. Some predictions are consistent with our previous deterministic models based on ODEs. For example, the *cln1*Δ *cln2*Δ *swi6*Δ *mbp1*Δ strain is predicted to be viable, whereas the *cln3*Δ *bck2*Δ *cdh1*Δ strain is inviable, which is consistent with corresponding predictions reported in our previous model (Kraikivski et al., 2015). Predictions that are consistent among different models are more reliable and can be confidently advanced for validation in wet labs.

Our modeling approach supplements the simplicity of Boolean models with quantitative details, such as real continuous time, and with easily interpretable, adjustable parameters that are helpful in accounting for mutant phenotypes. Therefore, our approach lies somewhere between Boolean models that lack quantitative details necessary to explain experimental observations and ODE models that provide all these details at the expense of estimating many obscure kinetic rate constants. In our approach, only the parameters that determine the delays must be estimated by fitting model simulations to experimental data. Also, our approach incorporates stochastic effects at minimal computational cost.

## Acknowledgments

K.T. acknowledges the Petchra Pra Jom Klao Ph.D. Research Scholarship (KMUTT – NSTDA) from King Mongkut’s University of Technology Thonburi (No: 103/2563).

## Data availability

All computer codes are available on Github: https://github.com/kittisaktaoma/A-continuous-time-Boolean-model-of-the-endocycle-events-in-budding-yeast

## Supplementary Information

**Supplementary Table S1:** Model parameter values.

| Supplementary Table S1: Model parameter values |  |  |  |
| --- | --- | --- | --- |
| A. Parameters defining the Boolean functions |  |  |  |
| Parameter | Value | Parameter | Value |
| $\omega_{Cdc14}$ | -0.5 | $\omega_{MBF}$ | -0.5 |
| $\omega_{Cdc14,Cdc5}$ | 1 | $\omega_{MBF,Clb2G}$ | 4 |
| $\omega_{Cdc14,Clb2M}$ | 1 | $\omega_{MBF,Clb5}$ | 3 |
| $\omega_{Cdc14,SAC}$ | 1 | $\omega_{MBF,Cln2}$ | 2 |
| $\omega_{Cdc14,SPOC}$ | 1 | $\omega_{MBF,Cln3}$ | 0.6 |
| $\omega_{Cdc20}$ | -0.5 | $\omega_{MBF,Nrm1}$ | 3 |
| $\omega_{Cdc20,Mcm1}$ | 1 | $\omega_{Mcm1}$ | -0.5 |
| $\omega_{Cdc20,SAC}$ | 1 | $\omega_{Mcm1,Clb2G}$ | 1 |
| $\omega_{Cdc5}$ | -0.5 | $\omega_{Mcm1,Clb2M}$ | 3 |
| $\omega_{Cdc5,Cdh1}$ | 2 | $\omega_{Nrm1}$ | 0.5 |
| $\omega_{Cdc5,Clb2G}$ | 3 | $\omega_{Nrm1,Cdh1}$ | 2 |
| $\omega_{Cdc5,Clb2M}$ | 3 | $\omega_{Nrm1,MBF}$ | 1 |
| $\omega_{Cdh1}$ | 0.5 | $\omega_{SBF}$ | -0.2 |
| $\omega_{Cdh1,Cdc14}$ | 3 | $\omega_{SBF,Clb2G}$ | 4 |
| $\omega_{Cdh1,Clb2G}$ | 1 | $\omega_{SBF,Clb5}$ | 1 |
| $\omega_{Cdh1,Clb2M}$ | 3 | $\omega_{SBF,Cln2}$ | 1 |
| $\omega_{Cdh1,Clb5}$ | 1 | $\omega_{SBF,Cln3}$ | 1 |
| $\omega_{Cdh1,Cln2}$ | 1 | $\omega_{SBF,Whi5}$ | 3 |
| $\omega_{Clb2G}$ | 0.5 | $\omega_{Sic1}$ | 1 |
| $\omega_{Clb2G,Cdh1}$ | 2 | $\omega_{Sic1,Clb2G}$ | 1 |
| $\omega_{Clb2G,Sic1}$ | 0.5 | $\omega_{Sic1,Clb2M}$ | 3 |
| $\omega_{Clb2M}$ | -0.5 | $\omega_{Sic1,Clb5}$ | 1 |
| $\omega_{Clb2M,Cdc20}$ | 2 | $\omega_{Sic1,Cln2}$ | 1 |
| $\omega_{Clb2M,Mcm1}$ | 1 | $\omega_{Sic1,Swi5}$ | 1 |
| $\omega_{Clb5}$ | -0.5 | $\omega_{Swi5}$ | -1 |
| $\omega_{Clb5,Cdc20}$ | 2 | $\omega_{Swi5,Cdc14}$ | 1 |
| $\omega_{Clb5,MBF}$ | 3 | $\omega_{Swi5,Mcm1}$ | 1 |
| $\omega_{Clb5,SBF}$ | 0.8 | $\omega_{Whi5}$ | 0.5 |
| $\omega_{Clb5,Sic1}$ | 0.25 | $\omega_{Whi5,Cdc14}$ | 1 |
| $\omega_{Cln2}$ | -1 | $\omega_{Whi5,Cln2}$ | 1 |
| $\omega_{\text{Cln2,MBF}}$ | 0.6 | $\omega_{\text{Whi5,Cln3}}$ | 1 |
| $\omega_{\text{Cln2,SBF}}$ | 1 | | |
| **B. Time constants and lognormal distributions** |  |  |  |
| **Parameter** | **Value** | **Parameter** | **Value** |
| $r_{\text{Glucose}}$ | $0.0077 \text{ min}^{-1}$ | $r_{\text{Galactose}}$ | $0.0046 \text{ min}^{-1}$ |
| $f_{\text{mean}}$ | 0.55 | $SPN_{\text{mean}}$ | 0.07 |
| $f_{\text{CV}}$ | 0.1 | $SPN_{\text{CV}}$ | 0.03 |
| $S_{0,\text{mean}}$ | 0.60 | $SPO_{\text{mean}}$ | 0.07 |
| $S_{0,\text{CV}}$ | 0.1 | $SPO_{\text{CV}}$ | 0.03 |

**Supplementary Table S2:** Reaction type classification.

| **Supplementary Table S2: Reaction type classification** |  |  |  |
| --- | --- | --- | --- |
| Reaction types | Gamma distribution | Mean (min) ± CV | Protein that is changing |
| Covalent modification | $k = 2, \theta = 0.3$ | $0.6 \pm 0.71$ | Whi5, MBF, SBF, Swi5, Cdh1, Cdc5, Mcm1 |
| Synthesis and degradation | $k = 3, \theta = 0.3$ | $0.9 \pm 0.58$ | Cln2, Clb5, Clb2, Cdc20, Cdc14, Nrm1, Sic1, Cln3 |
| No change | $k = 5, \theta = 0.3$ | $1.5 \pm 0.45$ | None |

**Supplementary Table S3:** Mutant strains simulated by the model. Inconsistencies between the model and experimental observations are indicated in red.

|  |  |  |  |  |  |
| --- | --- | --- | --- | --- | --- |
| Item | Mutant | Modified Parameter<br>(*i* any regulated proteins) | Phenotype |  |  |
|  |  |  | Experiment<br>(Reference) | ODE Model | Simulation |
| 1 | WT in glucose | $r = 0.0077$ | viable (size = 1.0)<br>(PMID: 6392855) | viable (size = 1.0)<br>(PMID: 27187804) | viable<br>(size = 0.54) |
| 2 | WT in galactose | $r = 0.0046$ | viable (size = 0.86)<br>(PMID: 32878900) | viable (size = 0.73)<br>(PMID: 27187804) | viable<br>(size = 0.44) |
| 3 | *whi5*Δ | $\omega_{i,Whi5} = 0$ | viable<br>(PMID: 12089449) | viable<br>(PMID: 28725464) | viable |
| 4 | *sic1*Δ | $\omega_{i,Sic1} = 0$ | viable<br>(PMID: 8614808) | viable<br>(PMID: 15169868) | viable |
| 5 | *swi4*Δ | $\omega_{i,SBF} = 0$ | viable<br>(PMID: 12024050) | viable<br>(PMID: 28725464) | viable |
| 6 | *mbp1*Δ | $\omega_{i,MBF} = 0$ | viable<br>(PMID: 8372350) | viable<br>(PMID: 28725464) | viable |
| 7 | *cdc5*Δ | $\omega_{i,Cdc5} = 0$ | inviable, no Cdc14<br>release<br>(PMID: 11832211) | inviable,<br>no Cdc14 release<br>(PMID: 28725464) | inviable<br>no Cdc14 release |
| 8 | *mcm1*Δ | $\omega_{i,Mcm1} = 0$ | inviable<br>(https://www.yeastgenome.org/locus/S000004646) | - | inviable |
| 9 | *cdc20Δ* | $\omega_{i,Cdc20} = 0$ | metaphase arrest<br>(*Cdc20* = 0 &<br>*Clb2M* = 1)<br>(PMID: 9501986) | inviable<br>(PMID: 15169868 &<br>PMID: 28725464 &<br>PMID: 27187804) | metaphase arrest<br>(*Cdc20* = 0 & *Clb2G* =<br>*Clb2M* = 1) |
| 10 | *cdc14Δ* | $\omega_{i,Cdc14} = 0$ | lethal-telophase<br>arrest<br>(*Cdc20* = 1 &<br>*Clb2G* = 1)<br>(PMID: 9885559) | telophase arrest<br>(*Cdc20* = 1 & *Clb2G* = 1<br>& *Clb2M* = 0)<br>PMID: 15169868 | telophase arrest<br>(*Cdc20* = 1 & *Clb2G* = 1<br>& *Clb2M* = 0) |
| 11 | *swi5Δ* | $\omega_{i,Swi5} = 0$ | viable<br>(PMID:12140549) | viable<br>(PMID: 27187804) | viable |
| 12 | *cdh1Δ* | $\omega_{i,Cdh1} = 0$ | viable<br>but smaller than wild<br>type<br>(PMID: 9288748) | viable<br>(PMID: 27187804 &<br>PMID: 15169868) | viable |
| 13 | *nrm1Δ* | $\omega_{i,Nrm1} = 0$ | viable<br>(PMID:12489128) | - | viable |
| 14 | Wild-type in<br>Nocodazole | SAC = 1 | metaphase arrest<br>(PMID: 10329618) | metaphase arrest<br>(PMID: 27187804) | metaphase arrest<br>(*Cdc20* = 0 & *Clb2G* =<br>*Clb2M* = 1) |
| 15 | *GAL-SIC1* | $\omega_{Sic1} = 1.5$<br>$r = 0.0046$ | viable<br>(PMID: 8164683) | viable<br>(PMID: 15169868) | viable |
| 16 | *GAL-CLN3* | $\omega_{Cln3} = 2$<br>$r = 0.0046$ | viable<br>(PMID: 1316273) | viable<br>(PMID: 27187804) | viable |
| 17 | *GAL-CLB5* | $\omega_{Clb5} = 0$<br>$r = 0.0046$ | viable<br>(PMID: 8319908) | viable<br>(PMID: 15169868) | viable |
| 18 | *GAL-CDC5* | $\omega_{Cdc5} = 1.5$<br>$r = 0.0046$ | viable<br>(PMID: 16455487) | viable<br>(PMID: 28725464) | viable |
| 19 | *GAL-CLB2* | $\omega_{Clb2G} = 2.5$,<br>$r = 0.0046$ | viable<br>(PMID: 8491189) | viable<br>(PMID: 27187804) | viable |
| 20 | *GAL-CDC20* | $\omega_{Cdc20} = 1.5$<br>$r = 0.0046$ | inviable<br>(PMID: 9482731) | inviable-mitotic<br>catastrophe<br>(PMID: 28725464) | inviable-mitotic<br>catastrophe<br>(*CDC20* = 1 before SPN<br>is finished) |
| 21 | *GAL-CDC14* | $\omega_{Cdc14} = 1.5$<br>$r = 0.0046$ | inviable (*G*<sub>1</sub> arrest)<br>(PMID: 9885559) | inviable (*G*<sub>1</sub> arrest)<br>(PMID: 27187804) | inviable-S arrest |
| 22 | *cln1Δ cln2Δ* | $\omega_{i,Cln2} = 0$ | viable<br>(PMID: 7588610) | viable<br>(PMID: 28725464) | viable |
| 23 | *cln3Δ bck2Δ* | $\omega_{i,Cln3} = 0$ | inviable (*G*<sub>1</sub> arrest)<br>(PMID: 10545447) | inviable (*G*<sub>1</sub> arrest)<br>(PMID: 15169868 &<br>PMID: 28725464) | inviable<br>(*G*<sub>1</sub> arrest: *Sic1* = 1,<br>*ORI* = 0) |
| 24 | *clb5Δ clb6Δ* | $\omega_{i,Clb5} = 0$ | viable<br>(PMID: 8319908) | viable<br>(PMID: 28725464) | viable |
| 25 | *clb1Δ clb2Δ* | $\omega_{i,Clb2G} = 0$<br>$\omega_{i,Clb2M} = 0$ | inviable (*G*<sub>2</sub> arrest)<br>(PMID: 1849457) | inviable (*G*<sub>2</sub> arrest)<br>(PMID: 28725464) | inviable (*G*<sub>2</sub> arrest)<br>(*Clb2G* = *Clb2M* = 0) |
| 26 | *cln3Δ bck2Δ*<br>*whi5Δ* | $\omega_{i,Cln3} = 0$<br>$\omega_{i,Whi5} = 0$ | viable<br>(PMID: 15210110) | viable<br>(PMID: 28725464) | inviable |
| 27 | *cln3Δ bck2Δ*<br>*sic1Δ* | $\omega_{i,Cln3} = 0$<br>$\omega_{i,Whi5} = 0$<br>$\omega_{i,Sic1} = 0$ | inviable<br>(PMID: 10545447) | inviable (*G*<sub>1</sub> arrest)<br>(PMID: 28725464) | inviable<br>(*G*<sub>1</sub> arrest: *Cdh1* = 1,<br>*ORI* = 0) |
| 28 | *cln1Δ cln2Δ*<br>*cdh1Δ* | $\omega_{i,Cdh1} = 0$<br>$\omega_{i,Cln2} = 0$ | viable<br>(PMID: 11809822) | viable<br>(PMID: 28725464) | viable |
| 29 | *cln1Δ cln2Δ*<br>*clb5Δ clb6Δ* | $\omega_{i,Cln2} = 0$<br>$\omega_{i,Clb5} = 0$ | inviable<br>(PMID: 7588610) | inviable (G<sub>1</sub> arrest)<br>(PMID: 15169868) | inviable (G<sub>1</sub> arrest)<br>(*Sic1* = *Cdh1* = 1) |
| 30 | *cln1Δ cln2Δ*<br>*sic1Δ* | $\omega_{i,Cln2} = 0$<br>$\omega_{i,Sic1} = 0$ | viable<br>(PMID: 7588610) | viable<br>(PMID: 15169868) | viable |
| 31 | *swi4Δ swi6Δ* | $\omega_{i,SBF} = 0$<br>$\omega_{i,MBF} = 0$ | inviable<br>(PMID: 8372350) | inviable (G<sub>1</sub> arrest)<br>(PMID: 28725464) | inviable |
| 32 | *swi5Δ cdh1Δ* | $\omega_{i,Swi5} = 0$<br>$\omega_{i,Cdh1} = 0$ | inviable (arrest as<br>binucleate cell)<br>(PMID: 12960422) | inviable<br>(telophase arrest)<br>(PMID: 15169868) | inviable<br>(telophase arrest)<br>(*CLB2G* = 1 and *ORI* = 1) |
| 33 | *cdc20Δ*<br>*clb5Δ clb6Δ* | $\omega_{i,Cdc20} = 0$<br>$\omega_{i,Clb5} = 0$ | inviable<br>(metaphase arrest)<br>(PMID: 10647015) | inviable<br>(metaphase arrest)<br>(PMID: 15169868) | metaphase arrest<br>(*Cdc20* = 0 & *Clb2G* =<br>*Clb2M* = 1) |
| 34 | *sic1Δ cdh1Δ* | $\omega_{i,Cdh1} = 0$<br>$\omega_{i,Sic1} = 0$ | inviable<br>(PMID: 9288748) | inviable<br>(PMID: 27187804) | inviable |
| 35 | *clb1Δ clb2Δ*<br>*clb5Δ clb6Δ* | $\omega_{i,Clb5} = 0$<br>$\omega_{i,Clb2G} = 0$<br>$\omega_{i,Clb2M} = 0$ | inviable (G<sub>1</sub> arrest) | inviable (G<sub>1</sub> arrest)<br>(PMID: 27187804) | inviable<br>(G<sub>1</sub> arrest:<br>*Sic1* = *Cdh1* = 1) |
| 36 | *GAL-SIC1*<br>*cln1Δ cln2Δ* | $\omega_{Sic1} = 1.5$<br>$\omega_{i,Cln2} = 0$<br>$r = 0.0046$ | inviable<br>(PMID: 11809822) | inviable (G<sub>1</sub> arrest)<br>(PMID: 15169868) | inviable |
| 37 | *GAL-SIC1 cln1Δ*<br>*cln2Δ cdh1Δ* | $\omega_{Sic1} = 1.5$<br>$\omega_{i,Cln2} = 0$<br>$\omega_{i,Cdh1} = 0$<br>$r = 0.0046$ | inviable<br>(PMID: 11809822) | inviable (G<sub>1</sub> arrest)<br>(PMID: 15169868) | inviable |
| 38 | *GAL-SIC1*<br>*swi5Δ cdh1Δ* | $\omega_{Sic1} = 1.5$<br>$\omega_{i,Cdh1} = 0$<br>$\omega_{i,Swi5} = 0$ | viable (rescued)<br>(PMID: 12960422) | viable<br>(PMID: 15169868) | viable |
| | | $r = 0.0046$ | | | |
| 39 | *GAL-CLB5 sic1Δ* | $\omega_{CLB5} = 0$<br>$\omega_{i,Sic1} = 0$<br>$r = 0.0046$ | inviable<br>(PMID: 10848575) | inviable<br>(ORI is not licensed)<br>(PMID: 15169868) | viable<br>(ORI is licensed) |
| 40 | *GAL-CLB5 cdh1Δ* | $\omega_{CLB5} = 0$<br>$\omega_{i,Cdh1} = 0$<br>$r = 0.0046$ | inviable<br>(PMID:12737808) | viable<br>(PMID: 15169868) | viable |
| 41 | *GALL-CDC20 sic1Δ cdh1Δ* | $\omega_{Cdc20} = 1.5$<br>$\omega_{i,Cdh1} = 0$<br>$\omega_{i,Sic1} = 0$<br>$r = 0.0046$ | viable<br>(PMID: 12737808) | viable<br>(PMID: 15169868) | inviable |
| 42 | *GAL-CLB2 cdh1Δ* | $\omega_{CLB2G} = 2.5$<br>$\omega_{i,Cdh1} = 0$<br>$r = 0.0046$ | inviable<br>(PMID: 12737808) | inviable<br>(telophase arrest)<br>(PMID: 15169868) | inviable<br>(telophase arrest)<br>*Clb2G* = 1 & *CLB2* = 0) |
| 43 | *GAL-CLB2 sic1Δ* | $\omega_{CLB2G} = 2.5$<br>$\omega_{i,Sic1} = 0$<br>$r = 0.0046$ | inviable<br>(telophase arrest)<br>(PMID: 9017392) | inviable<br>(PMID: 15169868) | inviable<br>(telophase arrest)<br>*Clb2G* = 1 & *CLB2* = 0) |
| 44 | *GAL-CLB2 swi5Δ* | $\omega_{CLB2G} = 2.5$<br>$\omega_{i,Swi5} = 0$<br>$r = 0.0046$ | inviable<br>(PMID: 9017392) | inviable<br>(telophase arrest)<br>(PMID: 15169868) | inviable<br>(telophase arrest)<br>*Clb2G* = 1 & *CLB2* = 0) |
| 45 | *swi4Δ swi6Δ sic1Δ* | $\omega_{i,SBF} = 0$<br>$\omega_{i,MBF} = 0$<br>$\omega_{i,Sic1} = 0$ | inviable<br>(PMID: 10545447) | inviable (*G*<sub>1</sub> arrest)<br>(PMID: 28725464) | inviable (*G*<sub>1</sub> arrest)<br>(*Cdh1* = 1) |
| 46 | *clb1-5Δ* | $\omega_{i,CLB2G} = 0, \omega_{i,CLB2M} = 0$<br>$\omega_{CLB5} = -1, \omega_{i,CLB5} / 8$ | *Clb6* oscillation<br>(PMID: 22306294) | *Clb6* oscillation<br>(PMID: 35504285) | *Clb6* oscillation |
| 47 | *GAL-CLB2-dbΔ* | $\omega_{\text{CLB2G}} = 1, \omega_{\text{CLB2M}} = 0,$<br>$r = 0.0046$ | Cdc14 oscillation<br>(PMID: 20660629) | Cdc14 oscillation<br>(PMID: 28725464 &<br>PMID: 35504285 &<br>PMID: 27187804) | Cdc14 oscillation |
| 48 | *GAL-CLB2-dbΔ*<br>*sic1Δ* | $\omega_{\text{CLB2G}} = 1, \omega_{\text{CLB2M}} = 0,$<br>$\omega_{i,\text{Sic1}} = 0$<br>$r = 0.0046$ | Cdc14 oscillation<br>(PMID: 20660629) | Cdc14 oscillation<br>(PMID: 28725464) | Cdc14 oscillation |
| 49 | *GAL-CLB2-dbΔ*<br>*swi5Δ* | $\omega_{\text{CLB2G}} = 1, \omega_{\text{CLB2M}} = 0$<br>$\omega_{i,\text{Swi5}} = 0$<br>$r = 0.0046$ | Cdc14 oscillation<br>(PMID: 20660629) | Cdc14 oscillation<br>(PMID: 28725464) | Cdc14 oscillation |
| 50 | *GAL-CLB2-dbΔ*<br>*cdc5Δ* | $\omega_{\text{CLB2G}} = 1, \omega_{\text{CLB2M}} = 0,$<br>$\omega_{i,\text{Cdc5}} = 0$<br>$r = 0.0046$ | no Cdc14 oscillation<br>(PMID: 20660629) | telophase arrest<br>Cdc14 is not released<br>(PMID: 28725464) | telophase arrest<br>(*Cdc14* = 0) |

**Supplementary Table S4:** Summary of time-delay parameters in endoreplication cycles and Cdc14 endocycles.

| Protein with time delay | Description | Gamma distribution (Mean ± CV) | |
| --- | --- | --- | --- |
|  |  | endoreplication | Cdc14 endocycle |
| Cdc5 | Delay for Cdc5 inactivation due to high Clb2G&M from GAL promotor | $0.6 \pm 0.7$ | $22.5 \pm 0.25$ |
| Cdh1 | Delay for Cdh1 inactivation when Clb5 is deleted | $22.5 \pm 0.25$ | $0.6 \pm 0.7$ |
| Cdh1 | Delay for Cdh1 activation due to high Clb2G&M level from GAL promotor | $0.6 \pm 0.7$ | $22.5 \pm 0.25$ |
| Clb6 | Delay for Clb6 activation because Clb6 has a lower transcription rate induced by MBF compared to Clb5 | $22.5 \pm 0.25$ | $0.9 \pm 0.5$ |

**Supplementary Figure S1.**
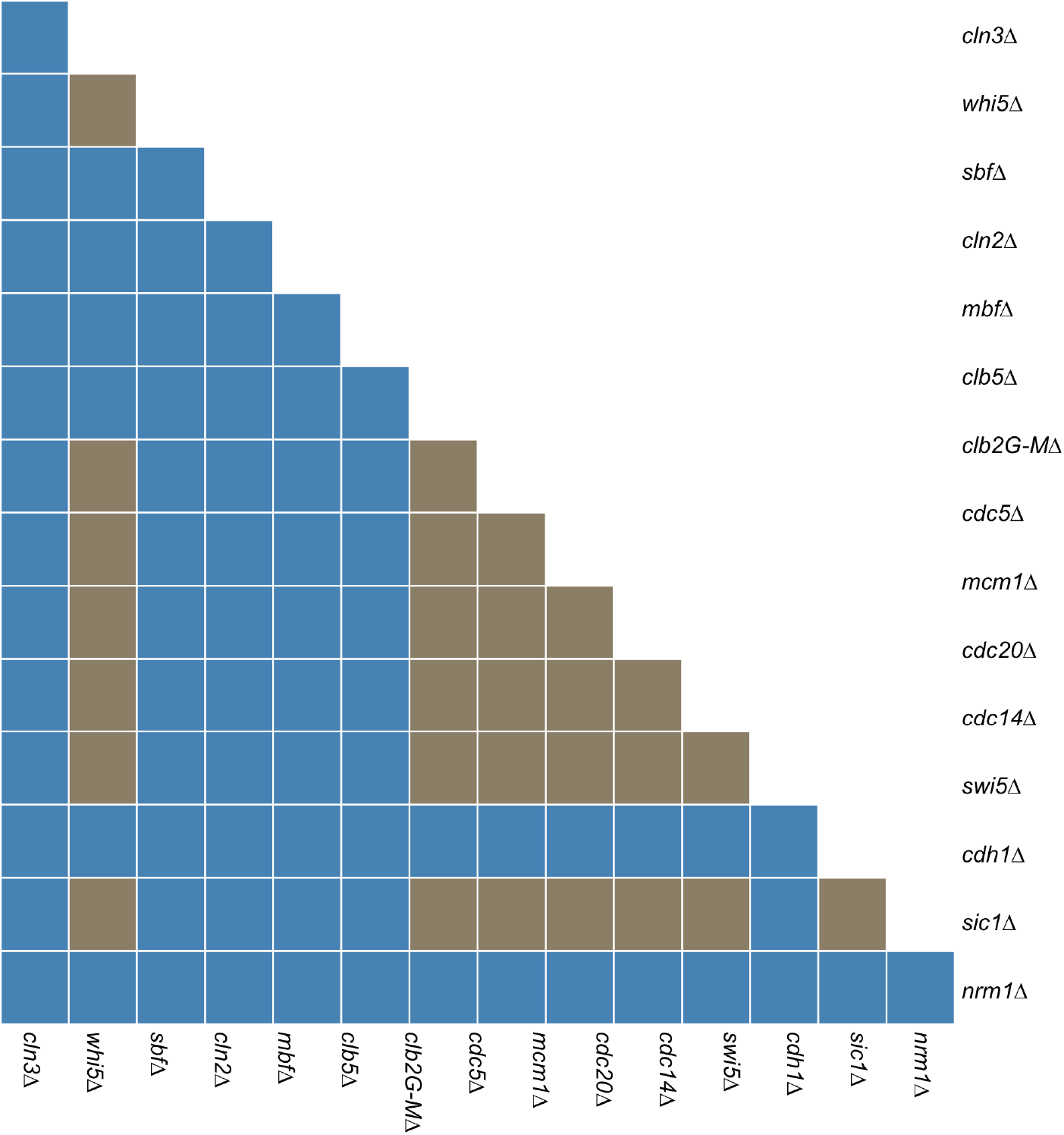
Predicted phenotypes when an additional deletion is added to the endoreplication strain (*clb1-5*Δ). (Blue) Mutants that lose Clb6 oscillations. (Brown) Mutants that retain Clb6 oscillations.

**Supplementary Figure S2.**
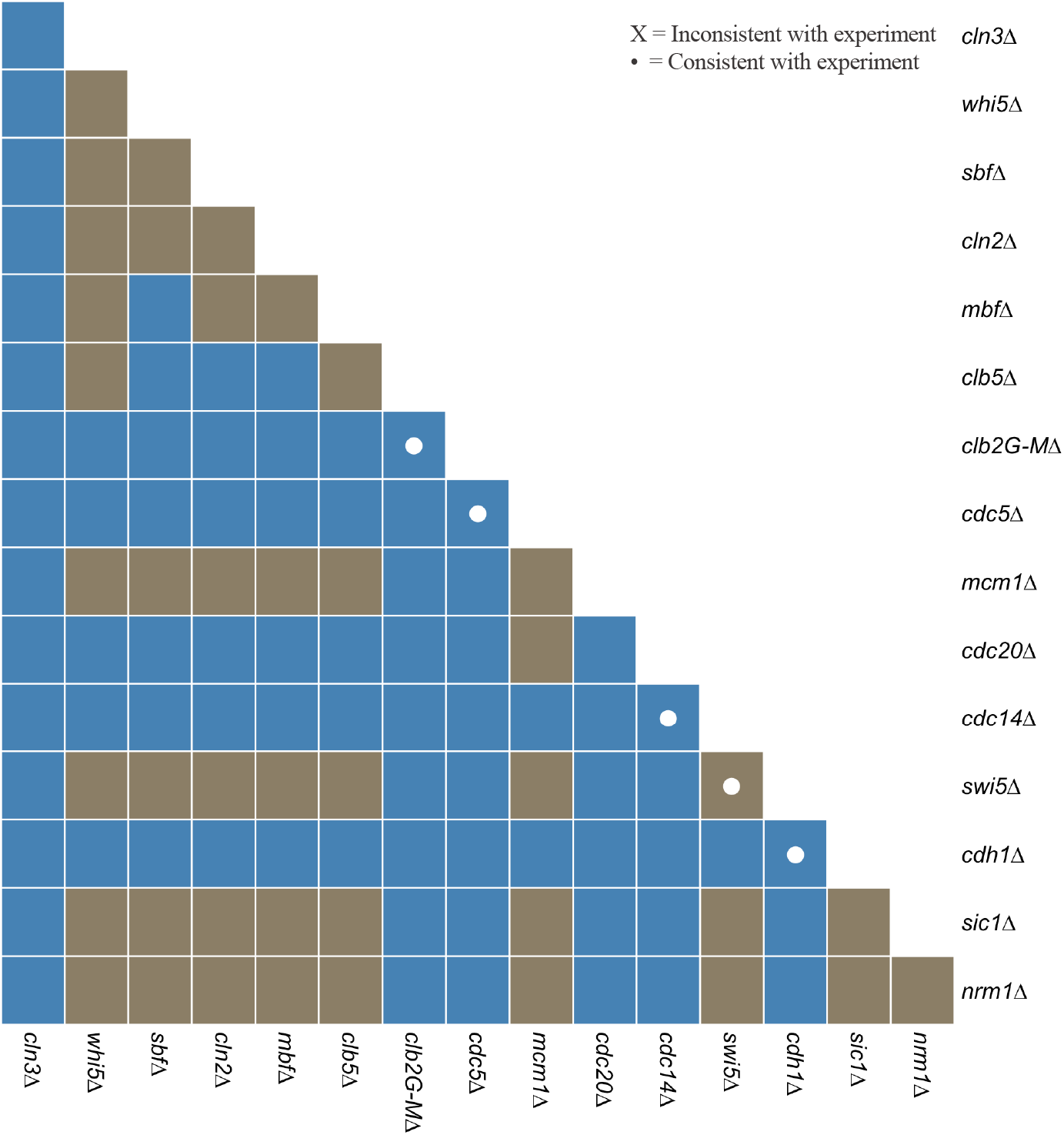
Predicted phenotypes when an additional deletion is added to the Cdc14 endocycle mutant (GAL*-CLB2-db*Δ). (Blue) Mutants that lose Cdc14 oscillations. (Brown) Mutants that retain Cdc14 oscillations. (White dot) Simulations that are consistent with experimental data.

